# One and Two Year Visual Outcomes from the Moorfields AMD Database - an Open Science Resource for the Study of Neovascular Age-related Macular Degeneration

**DOI:** 10.1101/450411

**Authors:** Katrin Fasler, Gabriella Moraes, Siegfried K. Wagner, Karsten U. Kortuem, Reena Chopra, Livia Faes, Gabriella Preston, Nikolas Pontikos, Dun Jack Fu, Praveen J. Patel, Adnan Tufail, Aaron Y. Lee, Konstantinos Balaskas, Pearse A. Keane

**Author notes:** **Correspondence and reprint requests:** Pearse A. Keane, MD FRCOphth, NIHR Biomedical Research Centre for Ophthalmology, Moorfields Eye Hospital NHS Foundation Trust and UCL Institute of Ophthalmology, United Kingdom. Tel: +44 Fax: +44.

## Abstract

**Objectives:** To analyse treatment outcomes and share clinical data from a large, single-center, well-curated database (8174 eyes / 6664 patients with 120,756 single entries) of patients with neovascular age related macular degeneration (AMD) treated with anti-vascular endothelial growth factor (VEGF). By making our depersonalised raw data openly available, we aim to stimulate further research in AMD, as well as setting a precedent for future work in this area.

**Setting:** Retrospective, comparative, non-randomised electronic medical record (EMR) database cohort study of the UK Moorfields AMD database with data extracted between 2008 and 2018.

**Participants:** 3357 eyes/patients (61% female). Extraction criteria were ≥ 1 ranibizumab or aflibercept injection, entry of “AMD” in the diagnosis field of the EMR, and a minimum of one year of follow-up. Exclusion criteria were unknown date of first injection and treatment outside of routine clinical care at Moorfields before the first recorded injection in the database.

**Main outcome measures:** Primary outcome measure was change in VA at one and two years from baseline as measured in Early Treatment Diabetic Retinopathy Study (ETDRS) letters. Secondary outcomes were the number of injections and predictive factors for VA gain.

**Results:** Mean VA gain at one-year and two years were +5.5±0.5 and +4.9±0.68 letters respectively. Fifty-four percent of eyes gained ≥5 letters at two years, 63% had stable VA (±≤14 letters), forty-four percent of eyes maintained good VA (≥70 letters). Patients received a mean of 7.7±0.06 injections during year one and 13.0±0.2 injections over two years.

Younger age, lower baseline VA, and more injections were associated with higher VA gain at two years.

**Conclusion:** This study benchmarks high quality EMR study results of real life AMD treatment and promotes open science in clinical AMD research by making the underlying data publicly available.

**Strengths and limitations of this study:**

- Large sample size, retrospective, single centre, electronic medical record database study
- High quality real life data
- Open science approach with sharing of depersonalised raw data

## INTRODUCTION

The treatment of neovascular age-related macular degeneration (AMD) has been revolutionised by the development of anti-vascular endothelial growth factor (VEGF) agents such as ranibizumab and aflibercept.(1–4) Unfortunately, real world results from retrospective studies are typically inferior to those from randomised controlled trials (RCTs), with fewer administered injections and significant inter-country and inter-center differences in therapy administration and outcomes.(5–9) Although retrospective studies and audits may be more likely than RCTs to reflect results in clinical practice, they still are not truly representative of outcomes in real world populations.(5–7,10,11) Major drawbacks of retrospective study designs are small sample sizes with selection bias and sub-optimal methods for handling of both missing data and losses to follow-up (LTFU).(11,12) Survival bias in particular can lead to skewed results: omission of cases LTFU from the analysis leads to selection of a non-random cohort with potential overestimation of visual acuity (VA) gains through exclusion of patients that stop treatment early due to irreversible visual loss such as foveal scarring or other adverse effects.

The advent of electronic medical records (EMR) has facilitated the collection of large amounts of data in routine clinical practice and thus has the potential to make retrospective study populations more representative of real life.(13–18) This is very much dependent, however, on the quality of data entry and the reliable follow-up of patients, and so these issues can remain problematic. The amount of data available from EMR systems also challenges the traditional methods of validation, analytics and reporting, and there is a struggle to implement the existing clinical research guidelines.(12, 19–21) For example, in 2015, the RECORD statement highlighted the challenges of using routinely collected observational health data.(21) A further problem is the variation of data collection in the different EMR registers in different hospitals and countries. The International Consortium for Health Outcomes Measurement (ICHOM) AMD working group has also proposed a standard set of clinical characteristics, interventions, and outcomes including preferential methods of VA recording (logarithm of the minimum angle of resolution or Early Treatment Diabetic Retinopathy Study (ETDRS) letters).(22)

At Moorfields Eye Hospital, an EMR was initiated in October 2008, and its successor, OpenEyes^TM^, was implemented in September 2012. Subsequently, data from both systems were merged into the current centralised repository, the data warehouse. We have created a dataset from this which represents, to our knowledge, the largest single-center cohort of patients receiving treatment for neovascular AMD in the world. This Moorfields AMD dataset is increasing steadily, with 909 new patients in 2017 alone, a number typically only comparable in magnitude to multicenter studies.(14,16) Apart from its sheer size, key advantages of this dataset include the ability to clean and validate data directly, the completeness due to the mandatory input of relevant fields including VA, the consistency of VA measurements in ETDRS letters, the lack of requirement to merge data from different sites and systems, the standardised treatment scheme following national guidelines, and the ability to directly access the raw imaging data from each study visit.

The aim of this study is to analyse one- and two-year VA outcomes, determine predictive factors of VA gain in treatment-naive eyes from the Moorfields AMD database, and to aid in scientific progress by making the de-personalised raw data from from our study openly available to the research community.(21,23)

## METHODS

### Data Collection

Data for this retrospective, comparative, non-randomised cohort study were extracted from the data warehouse, the centralised storage for all EMR data, of Moorfields Eye Hospital. Data were extracted between October 21, 2008 and August 08, 2018. Extraction criteria were ≥ 1 ranibizumab or aflibercept injection, entry of “AMD” in the diagnosis field of the EMR, and a minimum of one year of follow-up. Exclusion criteria were unknown date of first injection, any treatment outside of routine clinical care at Moorfields before the first recorded injection in the database, including pegaptanib, previous laser or photodynamic therapy, and bevacizumab. The rationale for exclusion of bevacizumab is that in the National Health Service (NHS), neovascular AMD is generally treated with the licensed therapeutics ranibizumab or aflibercept, and not with the off-label bevacizumab.(24,25) The date of the first injection was defined as the baseline date. The dataset has been depersonalised for publication and approval for data collection and analysis was obtained from the Institutional Review Board at Moorfields (ROAD17/031). The study adhered to the tenets set forth in the Declaration of Helsinki.

### Outcome Measures

The primary outcomes were mean change in VA from baseline as measured in ETDRS letters, proportion of eyes gaining ≥ 5 letters, proportion of eyes with stable vision (change in VA <15 letters to baseline), proportion of eyes with good vision (≥20/40 or 70 letters), and proportion of eyes with poor vision (≤20/200 or 35 letters). Those endpoints have been used in the pivotal trials and/or have been included in the ICHOM reporting recommendations.(1–4, 22) Secondary outcomes included the number of injections, and effect of baseline characteristics and injection numbers on changes in VA. Definitions for one-year and two-year outcome dates were taken from previous real-world studies as visits closest to 52 weeks and 104 weeks post baseline date within ±8 weeks.(6,13) We used the STROBE cohort checklist when writing our report.(20)

### Efforts to Minimize Bias

Clinical information from patients with neovascular AMD is manually entered to the Moorfields Eye Hospital EMR (OpenEyes^TM^) at each visit. The EMR requires mandatory completion for a number of fields at each patient visit, including VA, central retinal thickness, treatment decision, treatment drug, and injection number, thus minimizing the number of missing data entries. Of all 120,756 single entries, missing / zero visual acuity measurements were encountered in 4059 (4.1%) of all entries. After manual cleaning of all 4059 missing entries, missing data accounted for 808 (0.9%) entries. Patients aged <55 or >100 and eyes with injection numbers ≥50 were manually checked. Description of manual cleaning including a CONSORT diagram is shown in supplementary material (Supplementary 1, sFigure 1). Visual acuities below measurable ETDRS letters were converted to logMAR 2.0/-15 letters, logMAR 2.3/-30 letters and logMAR 2.7/-50 letters for count fingers, hand movements, and light perception respectively.(26) To avoid bias due to inter-eye correlation, statistical analysis was restricted to one eye per patient, i.e. the first eye of a patient if sequentially treated, and a randomly selected eye if simultaneously treated. Outcomes of second-treated fellow eyes will be reported separately.

### Statistical Analysis

The data were analysed using the statistics software R (https://www.r-project.org/; provided in the public domain by R Core team 2017 R Foundation for Statistical Computing, Vienna, Austria). The ggplot2 package was used for plots. The eye was defined as unit of analysis. Descriptive statistics included mean +/- 95% confidence interval (CI), and median, where appropriate. Differences between groups were evaluated using Mann Whitney U test and Pearson Chi-Square. Regression analysis was performed to assess relationship of predictive factors and VA gains. A *p* value of < 0.05 was interpreted as statistically significant.

### Patient and Public Involvement

Patients and public were not involved in the study as this was a retrospective cohort study.

### Data Sharing Statement

De-personalised data for this study will be openly available from the Dryad Digital Repository https://doi.org/10.5061/dryad.97r9289. Depersonalisation was carried out through hash function anonymisation of patient identification numbers, replacement of appointment dates with follow-up days to baseline, and categorising extreme age groups into age categories. Approval of adequate depersonalisation was obtained by Moorfields Information Governance.

## RESULTS

### Patient Demographics

The full dataset consisted of 8174 treatment-naïve eyes/6664 patients with 120,756 single entries treated for neovascular AMD in the Moorfields database between October 21, 2008 and August 9, 2018.

The dataset for analysis consisted of 3357 eyes/patients (61% female). Mean age was 78.2±0.3 years at baseline. Mean VA was 56.4±0.5 letters. Of these, 1105 eyes (33%) were treated with ranibizumab, 1533 (46%) with aflibercept, and 719 eyes (21%) were treated with both ranibizumab and aflibercept. The starting year of treatment ranged between 2007-2018. Therapeutic choices at Moorfields Eye Hospital changed after 2013 and both ranibizumab and aflibercept were offered as alternative first line agents. After this change, a number of patients were switched from one agent to another resulting in over 50% of eyes receiving both drugs during the course of treatment. Baseline characteristics are shown in Table 1.

**Table 1:**
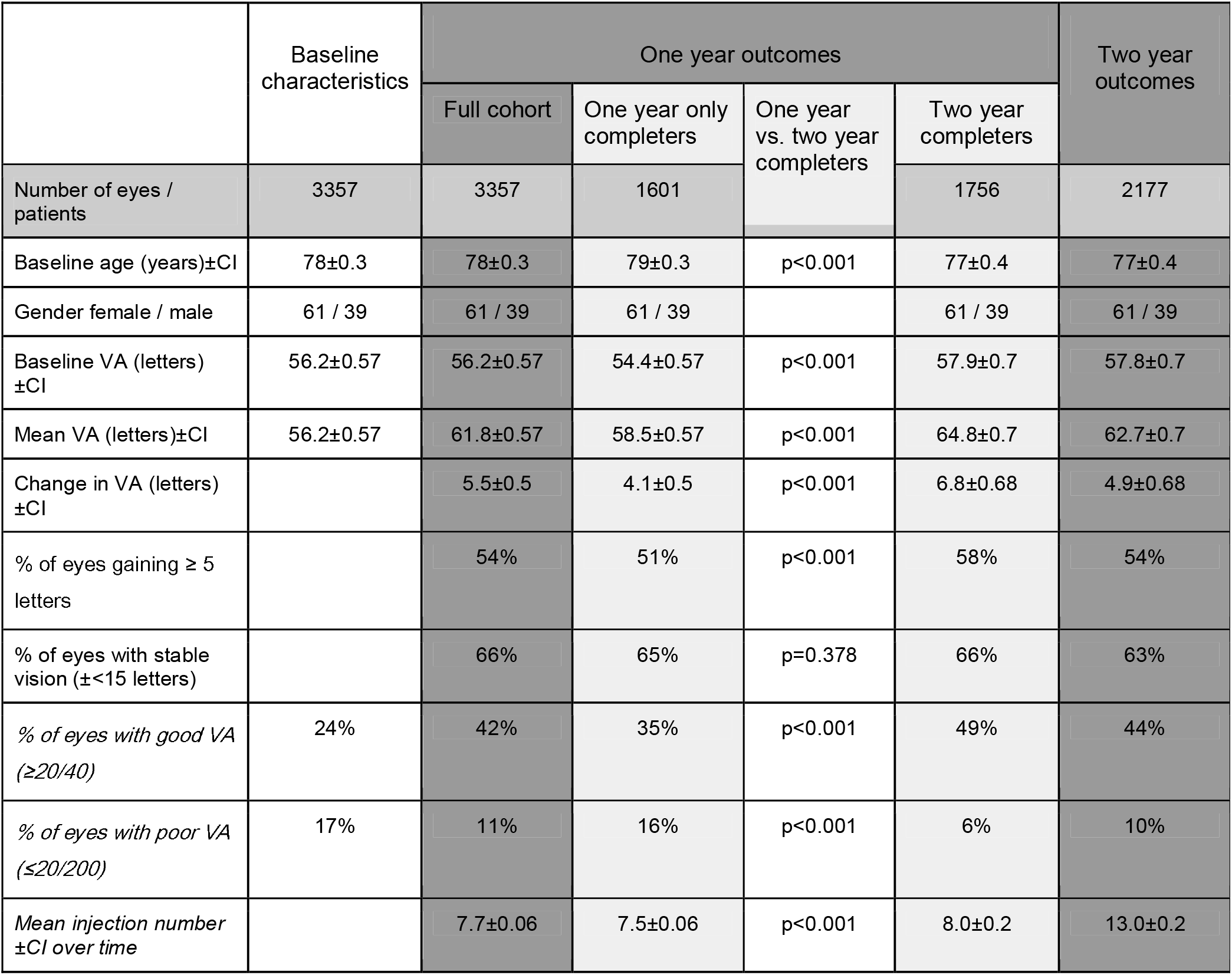
Baseline characteristics, one and two year outcomes, VA - visual acuity, CI - 95% confidence interval

Of the 1162 patients not completing the two year follow-up date, 254 patients had died. LTFU occurred in 27% of eyes for two year follow-up. To address the potentially resulting survival bias of, one-year outcomes for the cohort not completing 2-year follow-up and the cohort completing the 2-year follow-up are shown.

### Visual outcomes at one and two years

Mean VA gain at one and two years were +5.5±0.5 and +4.9±0.68 letters respectively. The mean number of injections over the first year and first two years were 7.7±0.06 and 13.0±0.2 respectively (Figure 1). Percentages of eyes gaining vision (change in VA ≥5 letters), stable vision (change in VA <15 letters), good vision (VA≥70 letters/>20/40), and poor vision (VA≤35 letters/≤20/200) are shown in Table 1 and Figure 2.

**Figure 1:**
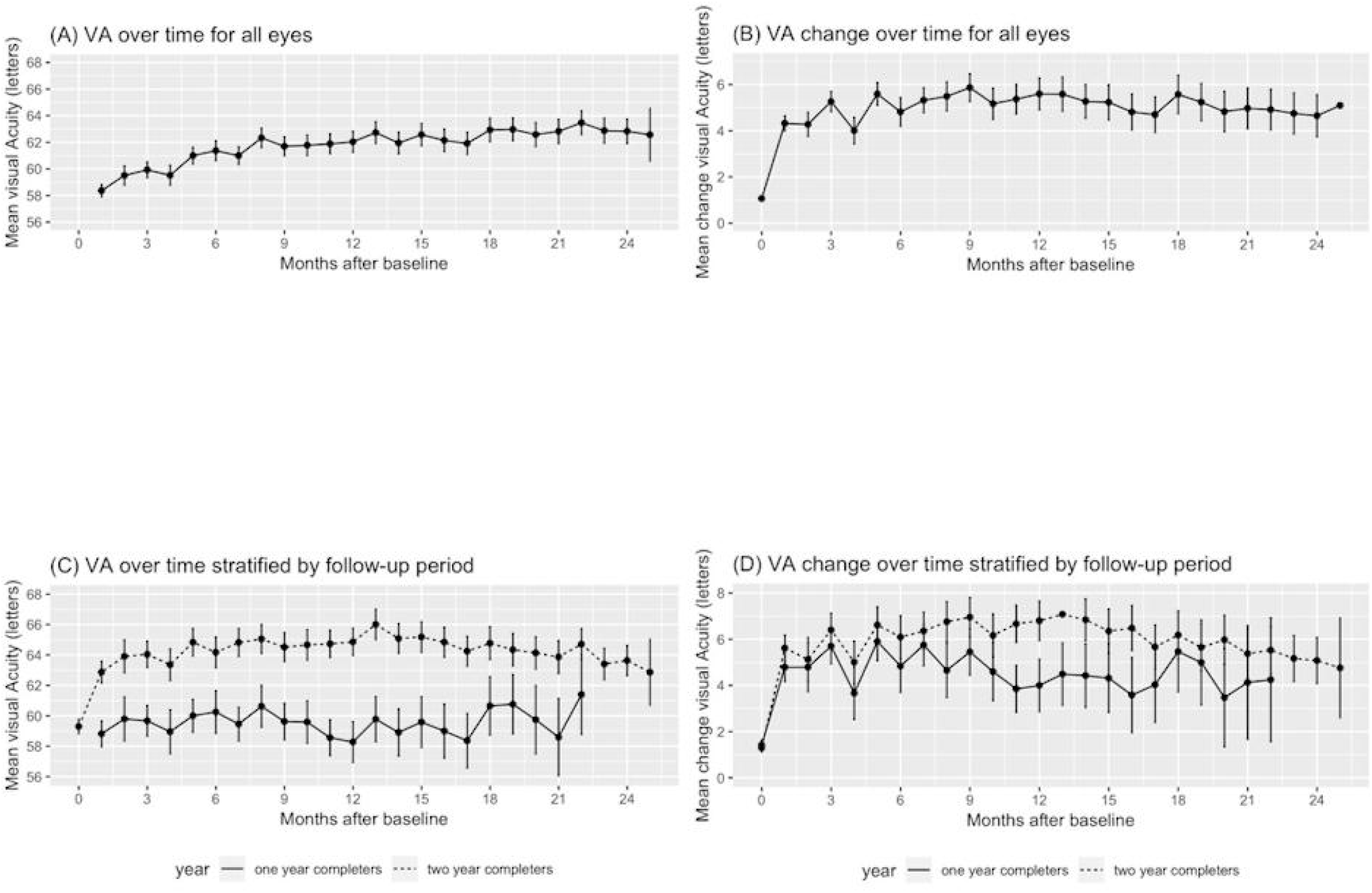
Visual acuity (A&C) and change in visual acuity (B&D) over time for all eyes and stratified by follow-up period (black: one year completers only; grey: two year completers). Bars represent 95% confidence intervals.

**Figure 2:**
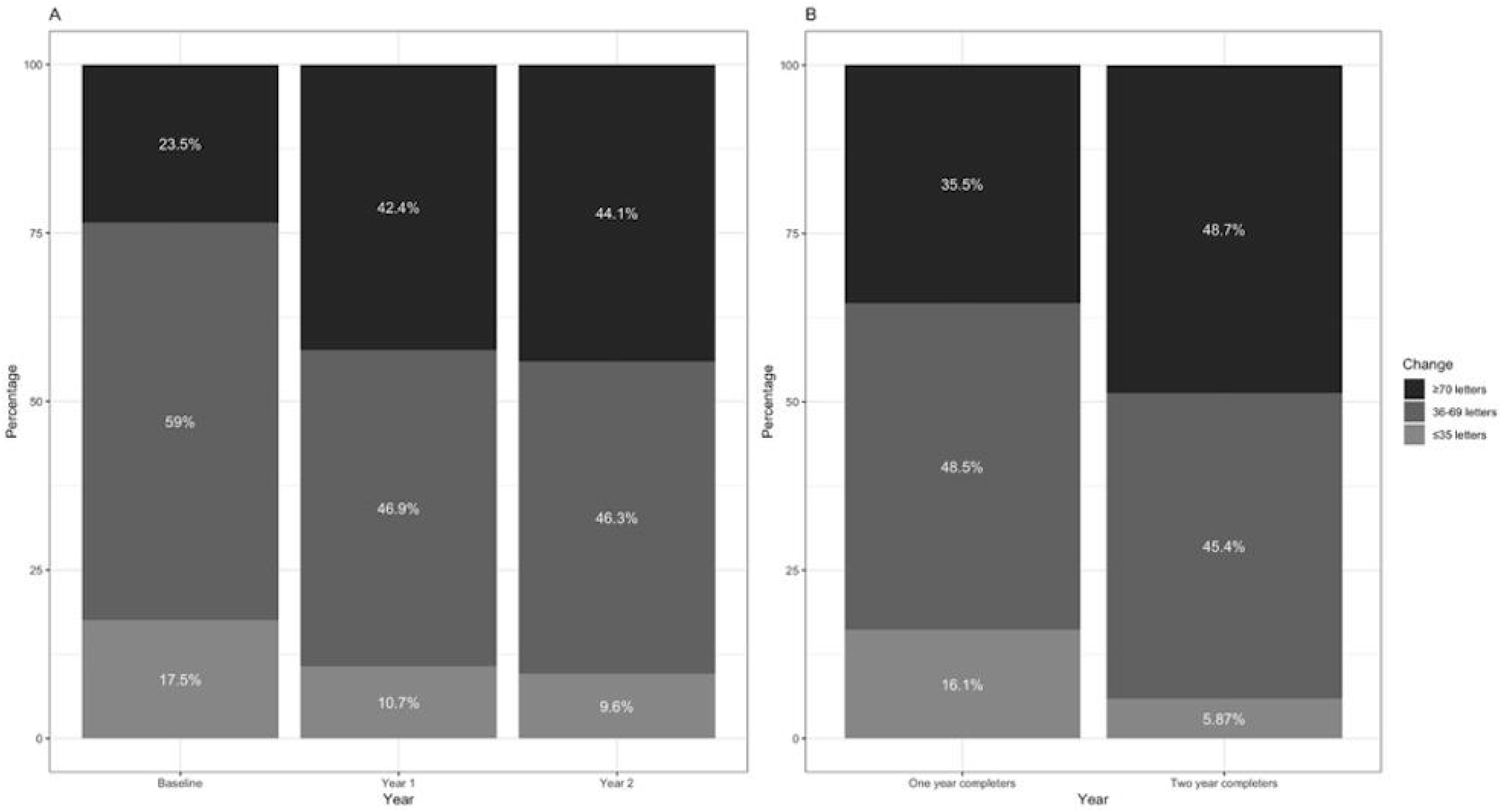
Percentage of eyes with good VA (≥ 70 letters), intermediate VA (36-69 letters), and poor VA (≤ 35 letters) at different follow-up times (A) and comparison of cohorts of different follow-up times at one year (B). VA - visual acuity

Comparison of subgroups that did not complete the two-year follow-up and the cohort that did complete the two-year follow-up showed a significantly lower mean baseline VA for those with a follow-up of less than two years (54.4 vs. 57.9 letters, p<0.05) a lower mean gain of letters (4.1 vs. 6.8 letters, p<0.05) as well as a lower injection frequency (7.5 vs. 8.0, p<0.05) at one year.

### Determinants of change in VA at one and two years

Regression to predict change in VA at one and two years from gender, baseline age, baseline VA, and injection number, showed only baseline VA, baseline age and injection number as significantly adding to the prediction (one year: F(8, 3384)=67.2, p<0.001, R^2^=0.1383); two years: F(8, 2168)=64.26, p<0.001, R^2^=0.1917). Lower baseline VA, lower baseline age, and higher number of injections are associated with a higher VA change at two years (Figure 3).

**Figure 3:**
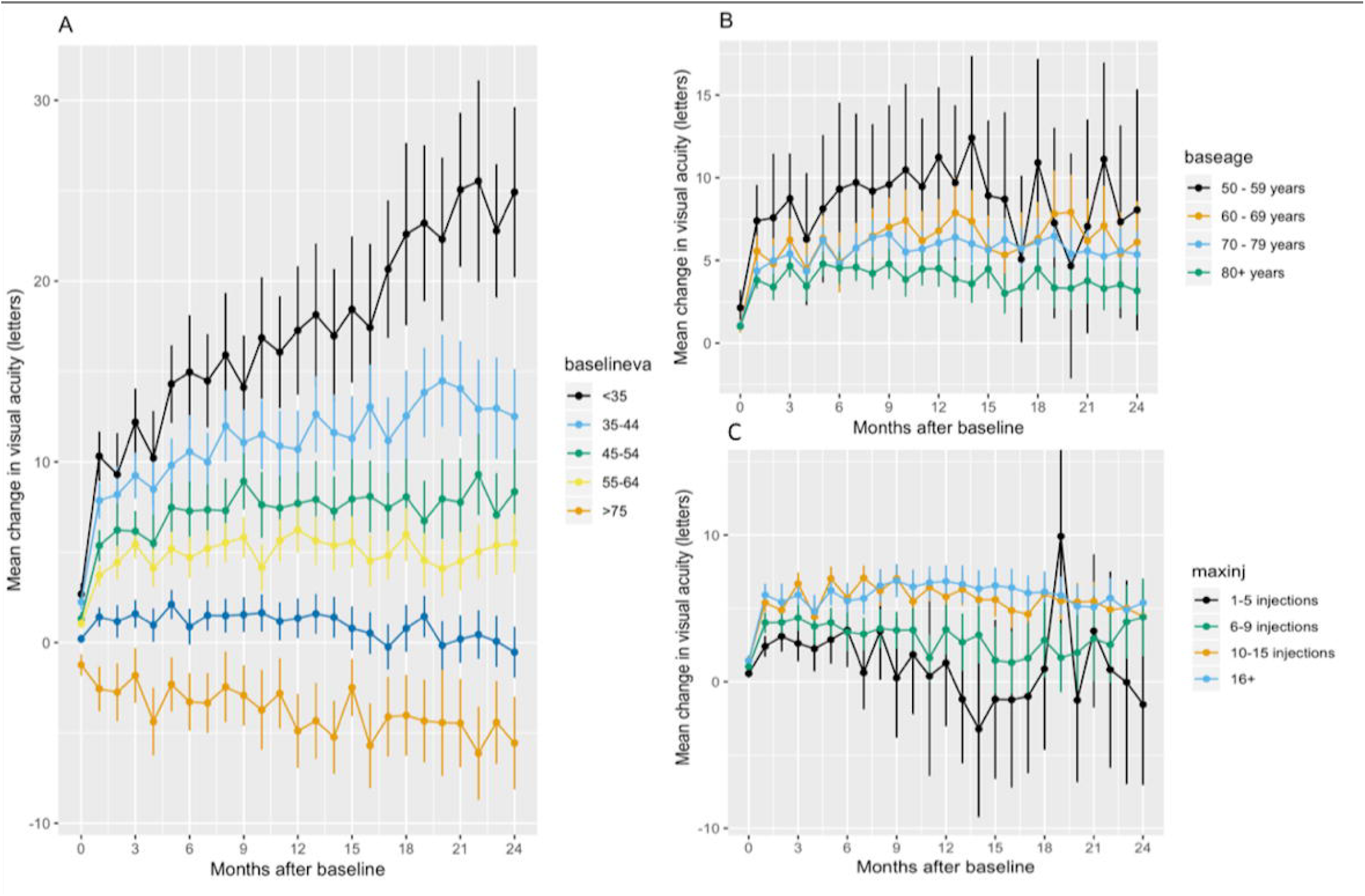
Change in visual acuity stratified by baseline VA (A), baseline age (B), and injection number at two years (C). Bars represent 95% confidence intervals.

## DISCUSSION

In this study, we show that patients treated with ranibizumab and/or aflibercept for neovascular AMD at Moorfields Eye Hospital achieve good visual outcomes, particularly those patients who present at an earlier age with better visual acuity, and who subsequently receive frequent intravitreal injections.

The Moorfields AMD Database is a large, consistent, and clean dataset of neovascular AMD treatment and visual outcomes, perhaps the largest single-center dataset of its kind worldwide. We have made this freely available to download with this manuscript in an effort to benefit the AMD research community. At a minimum this will allow for use of alternative statistical approaches and facilitate research reproducibility. (21) We also hope it will allow for the testing of new hypotheses and thus provide new insights into the treatment of this condition.(27) We have also developed systems so that the Moorfields AMD Database is automatically updated over time, with minimal need for manual cleaning of data. Just under 1000 new cases of neovascular AMD present to Moorfields Eye Hospital on a yearly basis - this may be particularly useful as new therapeutics for AMD continue to be introduced. Additionally, we plan on releasing data for long-term follow-up of these (five years and beyond), as well as their associated raw imaging data (colour fundus photography and optical coherence tomography (OCT) imaging in every eye at every visit).

At one and two years, our results of mean VA gains confirm the existing evidence in real-life studies, e.g., the Fight Retinal Blindness (FRB) group in Australia/New Zealand for ranibizumab/aflibercept with nearly identical baseline characteristics and visual acuity outcomes for mixed ranibizumab/aflibercept treatment (Table 2).(10,28)

**Table 2:** Comparison of two year outcomes with other real-life studies and randomised controlled trials. VA - visual acuity, A - aflibercept, R - ranibizumab, B - bevacizumab, EMR - electronic medical record, FRB - fight retinal blindness, CATT - comparison of Age-related Macular Degeneration Treatments Trials, VIEW - VEGF Trap-Eye: Investigation of Efficacy and Safety in Wet AMD

| Two year results | Retrospective, real-life studies |  |  |  | Prospective randomised trials |  |
| --- | --- | --- | --- | --- | --- | --- |
|  | Moorfields<br>R / A<br>1737 eyes | EMR Users<br>R<br>4990 eyes | FRB<br>B / R / A<br>1189 eyes | FRB<br>A<br>136 eyes | CATT<br>R / B<br>1107 eyes | VIEW<br>A<br>2063 eyes |
| Baseline age (years) | 77 | 80 | 79 | 77 | 79 | 75.6-76.5 |
| Baseline VA (letters) | 57.8 | 55 | 56.5 | 61.4 | 59.9-61.6 | 53.6-54.0 |
| Change in VA (letters) | 4.9 | +1 | +5.3 | +6 | 8.0-8.5 | 7.6-7.9 |
| % of eyes with good VA ($\geq 20/40$) | 44% | 30% | 45% | 58% | 67-68% | 30.7-34.9 |
| % of eyes with poor VA ($\leq 20/200$) | 10% | - | 11% | 10% | 4.7-8.4% | - |
| Mean injection number | 13.0 | 9.4 | 13 | 13.6 | 11.8 | 16.5 |

However, VA gains reported by the Writing Committee for the UK Age-Related Macular Degeneration EMR Users Group are considerably poorer which likely is explained by the reported capacity constraints resulting in reduced treatment frequency of with a mean of 9.4 injections over 2 years versus over 13 in our cohort and the FRB.(16) VA results from randomised prospective studies (e.g., the Comparison of Age-related macular degeneration Treatment Trials (CATT) and Vascular endothelial growth factor Trap-Eye: Investigation of Efficacy and Safety in Wet AMD study (VIEW)) have been shown to be superior to retrospective real-life data.(1,4)

This is also reflected in our data and is explained by the broader inclusion criteria, and the less strict treatment regimens with fewer administered injections. Comparison of cohorts that completed only one year of follow-up versus two or more years showed that eyes with shorter follow-up were older, had lower baseline VA, gained fewer letters at the one year follow-up, and received fewer injections over the first year. The loss to follow-up reflects the real-life setting of the study where patients transfer to stable AMD clinics, their vision has deteriorated and rendered further treatment unreasonable, or they are not able to further attend clinics. We deliberately did not perform any imputational replacement of missing data, but clearly describe the baseline characteristics and compare the one year results of the cohort LTFU before two years.(12)

VA gain over time is dependent on baseline characteristics and injection frequency.(12,14,29) Increasing age diminishes the VA gain expected as does a higher baseline acuity due to ceiling effect.(30) Baseline VA could even emerge as a surrogate measure for accessibility to treatment and quality of care, since simply looking at VA gains would underestimate centers that achieve short time from diagnosis to first treatment resulting in above average baseline VA but ceiling effect on VA gains.(8,12,16) Injection frequency has been recognised as another significant factor influencing VA gain and has been hypothesised to be the major factor in studies comparing ranibizumab and aflibercept due to the change in posology from treatment as needed to treat-and-extend concomitant with the change from ranibizumab to aflibercept in clinical practice.(14,29,31,32).

The retrospective nature and EMR-based data collection of our study introduce several limiting factors. Smoking status of our patients was not consistently available and thus, could not be included in the prediction model. Smoking has been identified as a risk factor for the development of neovascular AMD, but might also impact treatment response.(33) There is invariably survival bias within the data, as LTFU cannot be assumed to occur at random. However, baseline characteristics of LTFU as well as differences in outcomes for one and two year follow-up cohorts have been clearly described to address this. To date, there is no systematic collection of patient-reported outcome measures (mobility and independence, emotional well-being, as well as reading and accessing information questionnaires) as suggested by ICHOM.(22) The main advantages of this study are the quality and amount of data coming from one single center and one database. Moorfields Eye Hospital has a standardised treatment protocol for neovascular AMD, formerly treatment as needed, and fixed-first year/treat-and extend regimen with the introduction of aflibercept in 2014 (flow chart for aflibercept use is shown in Supplementary 2). The extensive manual cleaning and the homogeneous standards of data input (VA in ETDRS letters, mandatory fields) have formed a highly reliable resource which will be enhanced in the future with an automated update and validation to allow for continued growth and quality improvement of clinical AMD data.

In conclusion, this study shows that with a diligent approach, analysis of well maintained EMR data can lead to high quality real-life results and electronic availability of data facilitates maximisation of its potential in sharing research resources with the community, ultimately with the goal of improving patient care in real-life. In the near future, we plan to report on long-term visual outcomes (e.g., after 5-years), anatomic outcomes, and fellow-eye involvement, as well as the differential therapeutic effects of ranibizumab and aflibercept. In each case, we plan to release the raw data that underpins these reports - we hope that this will help promote an open-science approach to the study of neovascular AMD, and thus to direct patient benefit in the longer term.

## Author / contributorship statement

Katrin Fasler has drafted the manuscript and contributed to data acquisition, analysis, and interpretation of data. She is accountable for all aspects of the work and has approved for the final version to be published.

Siegfried K. Wagner and Karsten U. Kortuem have contributed in design of the study, interpretation of data as well as critical revision of the manuscript. They share accountability for all aspects of the work and have approved for the final version to be published.

Livia Faes, Dun Jack Fu, and Aaron Y. Lee have contributed to analysis and interpretation of the data and critical revision of the manuscript. They share accountability for all aspects of the work and have approved for the final version to be published.

Gabriella Moraes, Gabriella Preston, Reena Chopra, and Nikolas Pontikos have contributed to acquisition of data and critical revision of the manuscript. They share accountability for all aspects of the work and have approved for the final version to be published.

Konstantinos Balaskas, Praveen J. Patel, Adnan Tufail, and Pearse A. Keane have contributed to conception of the work, interpretation of data and critical revision of the manuscript. They share accountability for all aspects of the work and have approved for the final version to be published.

## Abbreviations

AMD: age related macular degeneration
VEGF: vascular endothelial growth factor
VA: visual acuity
ETDRS: Early Treatment Diabetic Retinopathy Study
RCT: randomized controlled trial
LTFU: lost to follow-up
EMR: electronic medical record
NHS: National Health Service
ICHOM: International Consortium for Health Outcomes Measurement FRB - Fight Retinal Blindness
CATT: Comparison of Age-related macular degeneration Treatment Trials
VIEW: Vascular endothelial growth factor Trap-Eye: Investigation of Efficacy and Safety in Wet AMD study

## Notes

**Disclosure / Funding** Dr. Fasler has received fellowship support from Alfred Vogt Stipendium and Schweizerischer Fonds zur Verhul□tung und Bekal□mpfung der Blindheit. She has been an external consultant for DeepMind. Dr. Wagner is an academic clinical fellow funded by the National Institute of Health Research. The views expressed in this publication are those of the author(s) and not necessarily those of the NHS, the National Institute for Health Research, Health Education England or the Department of Health. Dr. Keane has received speaker fees from Heidelberg Engineering, Topcon, Carl Zeiss Meditec, Haag-Streit, Allergan, Novartis, and Bayer. He has served on advisory boards for Novartis and Bayer and has been an external consultant for DeepMind and Optos. Dr. Keane is supported by a United Kingdom (UK) National Institute for Health Research (NIHR) Clinician Scientist Award (NIHR-CS--2014-12-023). The views expressed are those of the author and not necessarily those of the NHS, the NIHR or the Department of Health. Dr. Patel has received speaker fees from Novartis, UK, Bayer UK, and Roche UK and has received an advisory board honorarium from Novartis UK, Bayer UK. Dr. Lee has received research funding from Novartis, NVIDIA, and Microsoft Corporation. He is supported by the National Institute of Health (K23EY029246) and Research to Prevent Blindness.

